# Dysregulation of circular RNAs in myotonic dystrophy type 1

**DOI:** 10.1101/452391

**Authors:** Christine Voellenkle, Alessandra Perfetti, Matteo Carrara, Paola Fuschi, Laura Valentina Renna, Marialucia Longo, Rosanna Cardani, Rea Valaperta, Gabriella Silvestri, Ivano Legnini, Irene Bozzoni, Denis Furling, Carlo Gaetano, Germana Falcone, Giovanni Meola, Fabio Martelli

**Affiliations:** Molecular Cardiology Laboratory, IRCCS Policlinico San Donato, San Donato Milanese, Milan, Italy; Laboratory of Muscle Histopathology and Molecular Biology, IRCCS Policlinico San Donato, San Donato Milanese, Milan, Italy; Research Laboratories, IRCCS Policlinico San Donato, San Donato Milanese, Milan, Italy; Department of Geriatrics, Orthopaedic and Neuroscience, Institute of Neurology, Catholic University of Sacred Heart, Fondazione Policlinico Gemelli, Rome, Italy; Department of Biology and Biotechnology “Charles Darwin”, Sapienza University of Rome, Italy; Sorbonne Université, INSERM, Association Institut de Myologie, Centre de Recherche en Myologie, F-75013 Paris, France; Laboratory of Epigenetics, Istituti Clinici Scientifici Maugeri, Pavia, Italy; Institute of Cell Biology and Neurobiology, National Research Council, Monterotondo, Rome, Italy; Department of Neurology, IRCCS Policlinico San Donato, San Donato Milanese, Milan, Italy; Department of Biomedical Sciences for Health, University of Milan, Milan, Italy

**Author notes:** Corresponding author: Fabio Martelli, PhD, Molecular Cardiology Laboratory, IRCCS Policlinico San Donato, Via Morandi, 30-20097, San Donato Milanese, Milan, Italy.

**Keywords:** Musculoskeletal system, Muscular dystrophies, alternative splicing, circular RNA

## Abstract

Circular RNAs (circRNAs) constitute a recently re-discovered class of non-coding RNAs functioning as sponge for miRNAs and proteins, affecting RNA splicing and regulating transcription. CircRNAs are generated by “back-splicing”, linking covalently 3’- and 5’-ends of exons. Thus, circRNA levels might be deregulated in conditions associated to altered RNA-splicing. Indeed, increasing evidence indicates their role in human diseases. Specifically, myotonic dystrophy type 1 (DM1) is a multisystemic disorder caused by expanded CTG-repeats in the *DMPK* gene, resulting in abnormal mRNA-splicing. In this investigation, circRNAs expressed in DM1 skeletal muscles were identified by analyzing RNA-sequencing data-sets followed by qPCR validation. In muscle biopsies, out of 9 tested, 4 transcripts showed an increased circular fraction: CDYL, HIPK3, RTN4_03 and ZNF609. The circular fraction values correlated positively with skeletal muscle strength and Receiver-Operating-Characteristics curves showed that these four circRNAs allow to distinguish DM1 patients from controls. The identified circRNAs were also detectable in peripheral-blood-mononuclear-cells (PBMCs) and plasma of DM1 patients, but they were not regulated significantly, indicating a tissue-selectivity of the identified modulations. Finally, increased circular fractions of RTN4_03 and ZNF609 were also observed in differentiated myogenic cell lines derived from DM1 patients.

In conclusion, this proof-of-principle study identified circRNA dysregulation in DM1 patients.

## Introduction

Myotonic dystrophy type 1 (DM1), also known as Steinert disease (OMIM #160900), is an autosomal dominant multi-systemic disorder, with a spectrum of clinical manifestations that include myotonia, reduced muscle strength, cardiac arrhythmia, insulin resistance, cataracts, hypogonadism, and, in the most severe forms, cognitive defects [1-4]. The genetic defect in DM1 results from the dynamic expansion of CTG repeats in the 3’ untranslated region of the *dystrophia myotonica protein kinase (DMPK)* gene [5]. Severity of the disorder generally increases with the number of CTG repeats: healthy individuals have up to 40 repeats, patients with classic DM1 have 100-1000 repeats, and patients affected by congenital DM1 can have more than 2000 CTG repeats.

A major patho-mechanism underpinning DM1 is the generation of toxic RNAs containing expanded CUG triplets that accumulate as distinctive nuclear *foci and dysregulate the activity of* RNA processing factors, including MBNL1, CELF1, as well as Staufen1 and DDX5 [6-12]. Expanded CUG repeats have been demonstrated to be toxic *per se* in several cell types and animal models [13-15], disrupting pre-mRNA alternative splicing [16]. RNA splicing alterations result in the re-emergence of developmentally immature alternative splicing and polyadenylation patterns in adult muscles, as well as in alterations of localization and turnover of specific transcripts [7, 16-19].

Circular RNAs (circRNAs) are covalently closed loop-structure RNAs. They are generated by splicing events occurring on maturing pre-mRNAs in a different order than their genomic sequence, joining together a donor site with an upstream acceptor site [20-23].

CircRNAs have no accessible 5′borb3′ ends and are not poly-adenylated, escaping detection by many analytical and bioinformatics tools that are widely used in RNA biology. Indeed, for a long time, circRNAs were dismissed as rare aberrant by-products of the splicing process. More recently, large RNA deep-sequencing projects and the development of bioinformatics tools enabling the analysis of extensive data-sets, allowed the identification of significant proportions of back-splice junction reads associated to circRNAs in virtually all eukaryotic organisms [24-28]. While many circRNAs likely are byproducts of RNA splicing mechanisms, some circRNAs can be even more abundant than the linear counterparts [27] and specific biological functions have been associated to a rapidly increasing number of them. Certain circRNAs contain sequences complementary to the seed of a specific microRNA (miRNA), sequestering it and therefore reducing its bioavailability for target-mRNA inhibition [29]. A prototype of this is CDR1AS/ciRS-7, a circRNA containing about 70 evolutionarily conserved binding sites for miRNA-7 [25, 30]. Recently, CDR1AS was also found to regulate the turnover-rate of miR-7 in Cdr1as knock-out mice [31]. Certain circRNAs can regulate the expression of their linear counterparts, reducing the amount of pre-mRNA available for canonical splicing. Moreover, exon-intron circRNAs have also been described, that can interact with U1 snRNP and promote transcription of their parental genes [32]. Additionally, circRNAs binding and functionally interacting with RNA-binding proteins have been identified. For instance, circMBL, derived from the MBL/MBNL1 gene, in both D. melanogaster and humans, contains MBNL1 binding sites; MBL overexpression induces circMBL generation and this effect is dependent on the MBL binding sites [33]. Finally, a small fraction of circRNAs contains the necessary information to be translated with a cap-independent translation mechanism [26, 34].

While it is well established that RNA splicing is aberrant in DM1, whether circRNA levels are dysregulated has not been explored yet. With this study, we provide the first evidence that the levels of specific circRNAs linked to myogenesis are deregulated in skeletal muscle biopsies and in myogenic cell cultures derived from DM1 patients. Due to their resistance to exonucleolytic degradation, circRNAs are generally more stable than linear RNAs, constituting an attractive new class of potential biomarkers [35]. Accordingly, here we also show that circRNAs are detectable in both peripheral blood mononuclear cells (PBMCs) and plasma derived from the blood of DM1 patients.

## Methods

### RNAseq and Bioinformatics Analysis

Taking advantage of the GEO repository, we analysed RNAseq data-sets derived from *tibialis anterior* muscle biopsies, taken from DM1 and control patients (GSE86356) [36]. In order to ensure sufficient sequencing depth for the identification of the expectedly rare back-splicing events, we used only data-sets containing more than 75 million reads (Table S1). In this way, raw reads in fastq format from 5 control and 25 DM1 ribo-depleted libraries were aligned to hg19 reference genome with BWA software (v. 0.7.12), choosing the options according to CIRI2 manual [37, 38].

Subsequently, circRNAs were identified by detecting back-splice events in each of the aligned samples using CIRI2 (v. 2.0.6) with the suggested parameters. All identified circRNAs were then collected, normalized to each library size and quantified using custom R scripts. An abundance filter was applied by removing back-splice events present in <70% of either control or DM1 samples (Table S2). The circular-to-linear ratio was calculated using the highest expressed linear junction involved in the relevant back-splice event (Table S3). Briefly, we compared all annotated linear junctions that comprised either the acceptor or the donor site of the back-splice junction and kept the one with the highest amount of spliced reads. This ensured that the ratio was determined by using two equal biological entities, i.e. reads spanning a splice junction.

### Patient characteristics and tissue collection

The clinical diagnosis of DM patients was based upon the criteria set by International Consortium for Myotonic Dystrophies guidelines [39]. Genetic analysis was carried out to confirm DM1 diagnosis as described previously [40]. The Muscular Impairment Rating Scale (MIRS) was used to determine the disease stage [41]. MRC scale (Medical Research Council) was used to evaluate muscle strength.

*Biceps brachii* muscle biopsies collected from 20 DM1 patients and 19 sex-and age matched subjects without signs of neuromuscular disorders (controls) were used for validation (Table 1).

**Table 1.**
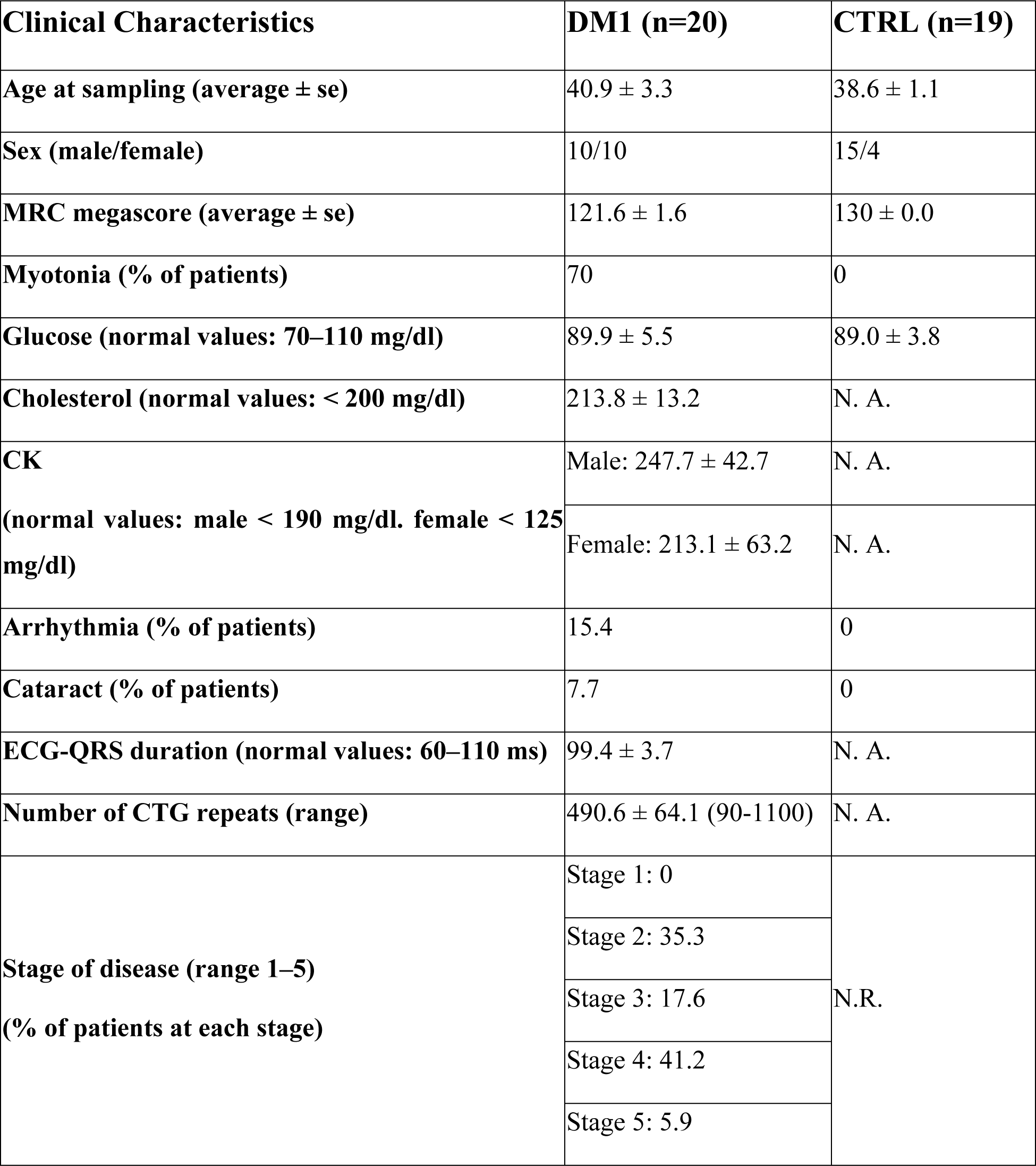
Clinical data on DM1 and control patients used for validation in biopsies. N.R.: not relevant. N.A.: not available.

PBMCs were isolated from the peripheral blood of 19 DM1 and 18 sex- and age matched controls (Table S4) by Ficoll-Paque™ PLUS (Ge Healthcare) gradient centrifugation as described before [42]. The plasma of 29 DM1 and 28 age-and sex-matched controls (Table S4) was collected in EDTA-tubes and cells as well as platelets were removed as described previously [43, 44].

### Ethical approval and informed consent

The experimental protocol was reviewed and approved by the Institutional Ethics Committee of the San Raffaele Hospital (protocol miRNADM of 23.06.2015) and was conducted according to the principles expressed in the Declaration of Helsinki, the institutional regulation and the Italian laws and guidelines. A written informed consent was obtained from each patient prior to muscle biopsies or blood collection.

### Histopathological analysis

Muscle tissue was fresh-frozen in isopentane cooled in liquid nitrogen. Histopathological analyses were performed on serial sections (8 µm) processed for routine histological or histochemical stainings. Myofibrillar ATPase staining was performed as previously described, after sample pre-incubation at pH 4.3, 4.6, and 10.4 [45].

### DM1 myogenic cell lines

Immortalized human myotonic dystrophy muscle cell lines expressing murine *Myod1* cDNA under the control of a Tet-on inducible construct were previously described [46]. Cells were cultivated in DMEM growth medium (15% FBS) until confluency was reached. Myogenic differentiation was induced by switching cell cultures to DMEM supplemented with 5 µg/ml insulin and 4 µg/ml doxycycline (Sigma-Aldrich).

For silencing experiments, cells were transfected after 3 days of differentiation with 50nM MBNL1/CELF1 TARGETplus SMARTpool siRNAs (Dharmacon) or with 50nM ON-TARGETplus Non-targeting Pool as negative control. Cells were transfected using HiPerFect reagent (Qiagen), according to manufacturer’s instructions. Three days after transfection, total RNA was isolated and analyzed by qPCR.

### Isolation of total RNA

Total RNA was extracted from the muscle tissues using TRIzol reagent (Thermo Fisher Scientific Inc.) as described previously [47, 48]. For isolation of total RNA from PBMCs and cells, TRIzol reagent was used according to the manufacturer’s instructions. The purity and integrity of the obtained RNAs were measured by Nanodrop (Thermo Fisher Scientific Inc.).

Total RNA from plasma samples was extracted as previously described using NucleoSpin miRNA Plasma columns (Macherey-Nagel) [43, 44].

### Real-time reverse transcriptase qPCR

For validation experiments, total RNA was first retro-transcribed using the SuperScript Reverse Transcriptase kit (version III or IV) and then investigated by SYBR green qPCR, according to the manufacturer’s protocol (Thermo Fisher Scientific Inc.). Primer couples were designed by Primer-BLAST tool (Table S5).

The relative expression was calculated using the comparative Ct method 2^−⋄⋄CT^ [49], normalizing to the averaged Cts of RPL13, RPL23 and UBC for tissue RNAs and to the averaged Cts of miR-106a and miR-17-5p for plasma RNAs [43, 44].

The circular-to-linear ratio was estimated by subtracting the raw Ct of the linear transcript from the raw Ct of the corresponding circular transcript.

For score-calculations, the log2 fold changes of all significantly modulated circRNAs or circular-to-linear ratios were averaged.

### Statistical Analysis

GraphPad Prism 7.01 (GraphPad Software Inc.) was used for statistical analysis and for graph generation. Following differential expression analysis, all data-sets were checked for their distribution by D’Agostino & Pearson normality test. The only exception were the cell line data, since due to the small sample size normality test was not possible. The differential expression of circRNAs in biopsies was investigated by multiple t-test, using the recommended settings of GraphPad for false discovery rate (Benjamini, Krieger and Yekutieli) and Q= 1% as significance cut-off. For differential expression analysis of circRNAs in PBMCs, plasma and cell lines, two-tailed Student’s t test or Mann Whitney was used, depending on the data-distribution. A p <0.05 was deemed statistically significant. Values are expressed as ±standard error.

## Results

### Identification of circRNA expression in DM1 skeletal muscle by RNA-sequencing

Published RNA sequencing (RNAseq) data of ribo-depleted libraries derived from 5 controls and 25 DM1 *tibialis anterior biopsies [36] were investigated for circRNA expression. Among the* available data-sets, we analyzed only libraries containing at least 76 million reads, thus providing sequence information at a high depth (Table S1). By applying CIRI2 [37, 38], an algorithm designed for the discovery of circRNAs, a total of 21.822 unique back-splice sites were identified across all libraries. Since most of these events were present in few samples only, a stringent abundance filter was used, resulting in ≈b.0éyybback9splicebjunctionsbD±ableb™H60b:ertainbcirc%WTsbdisplayb expression levels comparable to or even higher than their linear counterparts, suggesting a potential biological relevance [27]. To identify these particularly interesting circRNAs, the circular-to-linear ratios were estimated. Therefore, all annotated linear junctions involved with either the donor or the acceptor site of the back-splice event were quantified. The linear junction with the highest expression was then compared to the back-splice junction, revealing 578 circRNAs with a circular-to-linear ratio >0.5 in DM1 (Table S3).

To narrow the list of candidates for validation, the ≈ 1.800 identified circRNAs were intersected with a list of 29 circRNAs that were previously validated in human and mouse myoblasts [26], resulting in 18 common circRNAs (Table S3). Interestingly, most of them displayed a circular-to-linear ratio >0.5.

### Validation by qPCR of differentially expressed circRNAs in DM1 skeletal muscles

Muscle tissue biopsies were harvested from *biceps brachii* of 20 DM1 and 19 sex- and age-matched control individuals, with no sign of neuromuscular disorders. The DM1 group showed the main characteristics of the disease, such as myotonia, cataract and muscle weakness (Table 1), as well the typical histological alterations, such as central *nuclei*, high variation in fiber size, atrophic fibers and nuclear clumps (Fig. S1) [1, 39]. Most DM1 patients were at stage 2-4 and the pathological expansions of the CTG triplets ranged from 90 to 1100.

Total RNAs were isolated and the expression of a set of circRNAs and their linear counterparts was measured by qPCR. Out of the 18 circRNA candidates, 8 primer couples passed all technical checks of specificity and efficiency. Of note, two circRNAs originated from the same gene and were indicated as circRTN4 and circRTN4_03. The primers designed for circRNAs produced an amplicon spanning the back-splice junction, while the linear primers resulted in amplicons crossing the linear junction to a neighboring exon. Due to the key role of MBNL in DM1, the previously identified circRNA hosted in the second exon of MBNL [33] and its linear form were also measured. Thus, a set of 9 circRNAs together with their linear counterparts were used for validation (Table S4). All tested transcripts were confirmed to be readily expressed also in *biceps brachii* biopsies, with the only exception of circMBNL1 (circMBNL1), showing an expression close to the detection threshold. Five circRNAs, circASPH, circCDYL, circHIPK3, circRTN4_03 and circZNF609, displayed a statistically significant increase following multiple comparison testing (q<0.01) (Fig. S2).

To assess whether the observed induction of the circular transcripts was simply the consequence of a general increase of transcription in the relevant genomic region in DM1 patients, modulation of the ratios between the circular and the linear isoforms was calculated. We identified 4 circRNAs (circCDYL, circHIPK3, circRTN4_03 and circZNF609) with a significantly increased, circular-to-linear ratio in DM1 muscles (Fig. 1a), implying a de-regulation of the circular transcript independent from its linear counterpart. Accordingly, a similar trend was also observed in the RNAseq data, where the circular-to-linear ratios were higher in the DM1 affected muscles compared to the controls (Table S3).

**Figure 1.**
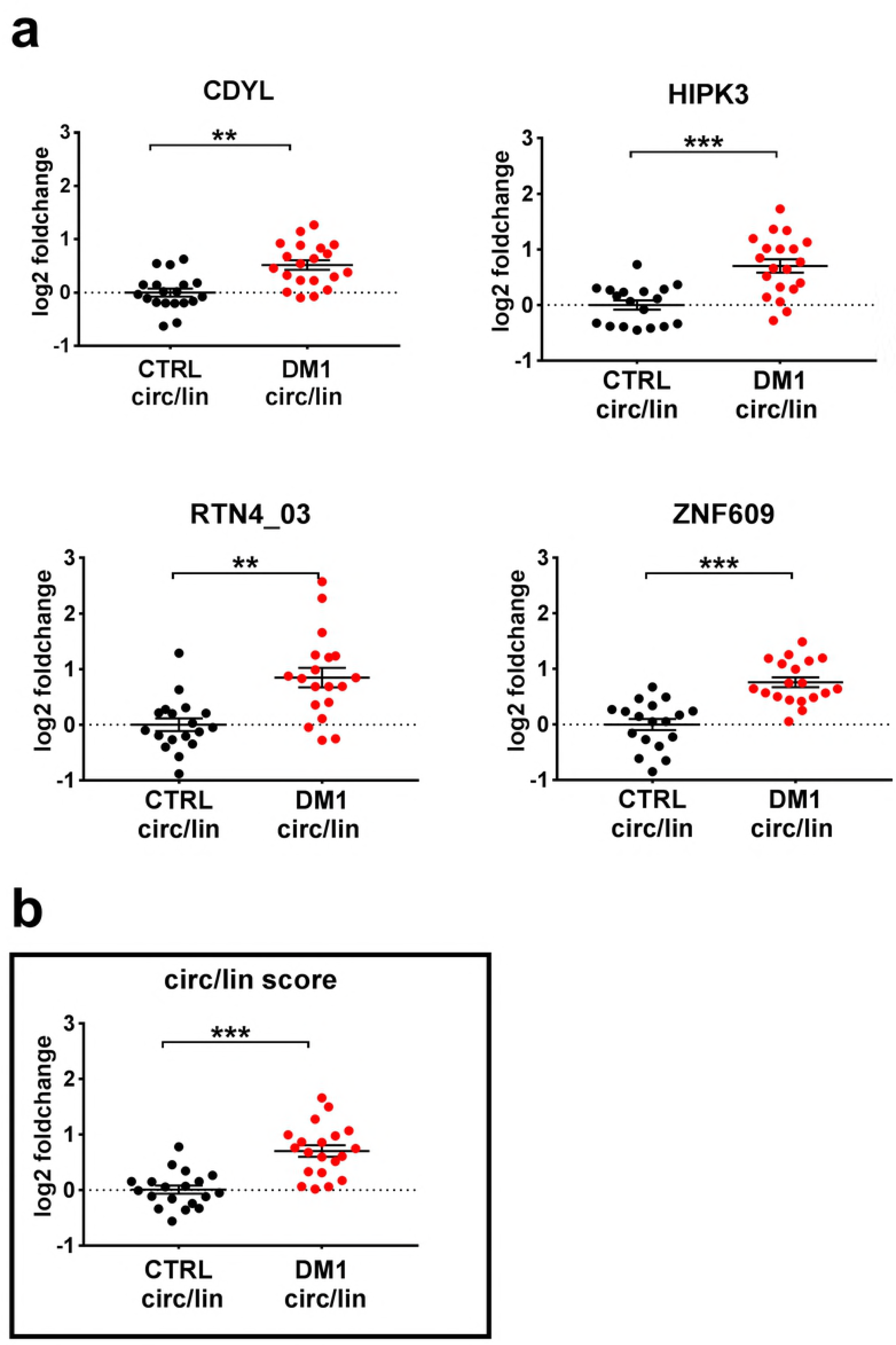
Differentially modulated circular-to-linear ratios in DM1 skeletal muscles. **(a)** Scatterplots in log2 scale of significantly different circular-to-linear ratios identified by qPCR in DM1 muscle tissue compared to control (CTRL). After normality test, statistical significance was calculated either by t-test or Mann-Whitney test (threshold p<0.05), followed by correction multiple comparison, with significance threshold set at q<0.01. **(b)**Scatterplot of circular-to-linear ratio, estimated by averaging the log2 fold changes of significantly different circular-to-linear ratios in DM1 muscle tissue compared to control. For both panels, lines indicate mean and standard error values for each group. DM1= 20 (red dots); CTRL= 19 (black dots); **q<0.001; ***p<0.001.

### DM1-circRNAs distinguish DM1 patients from controls

To understand if the identified DM1-deregulated circRNAs (DM1-circRNAs) display a discriminating power to identify DM1 patients, Receiver Operating Characteristic (ROC) curve analysis was performed. Both, significantly increased circRNAs alone (Fig. S3) and circular-to-linear ratios (Fig. 2) were analyzed. Interestingly, with one exception (circCDYL), the discrimination power between diseased and healthy individuals increased using the circular-to-linear ratios. In detail, among the five tested ratios, ZNF609 showed the largest area under the curve (AUC= 0.92), while the others ranged between 0.84 and 0.86 (Fig. 2a). Intriguingly, averaging all five DM1-circRNA fractions into a “circular-to-linear score” (Fig. 2b) improved the performance (AUC= 0.89) with respect to the singular fractions, with the exception of circZNF609, continuing to show the largest AUC (Fig. 2a).

**Figure 2.**
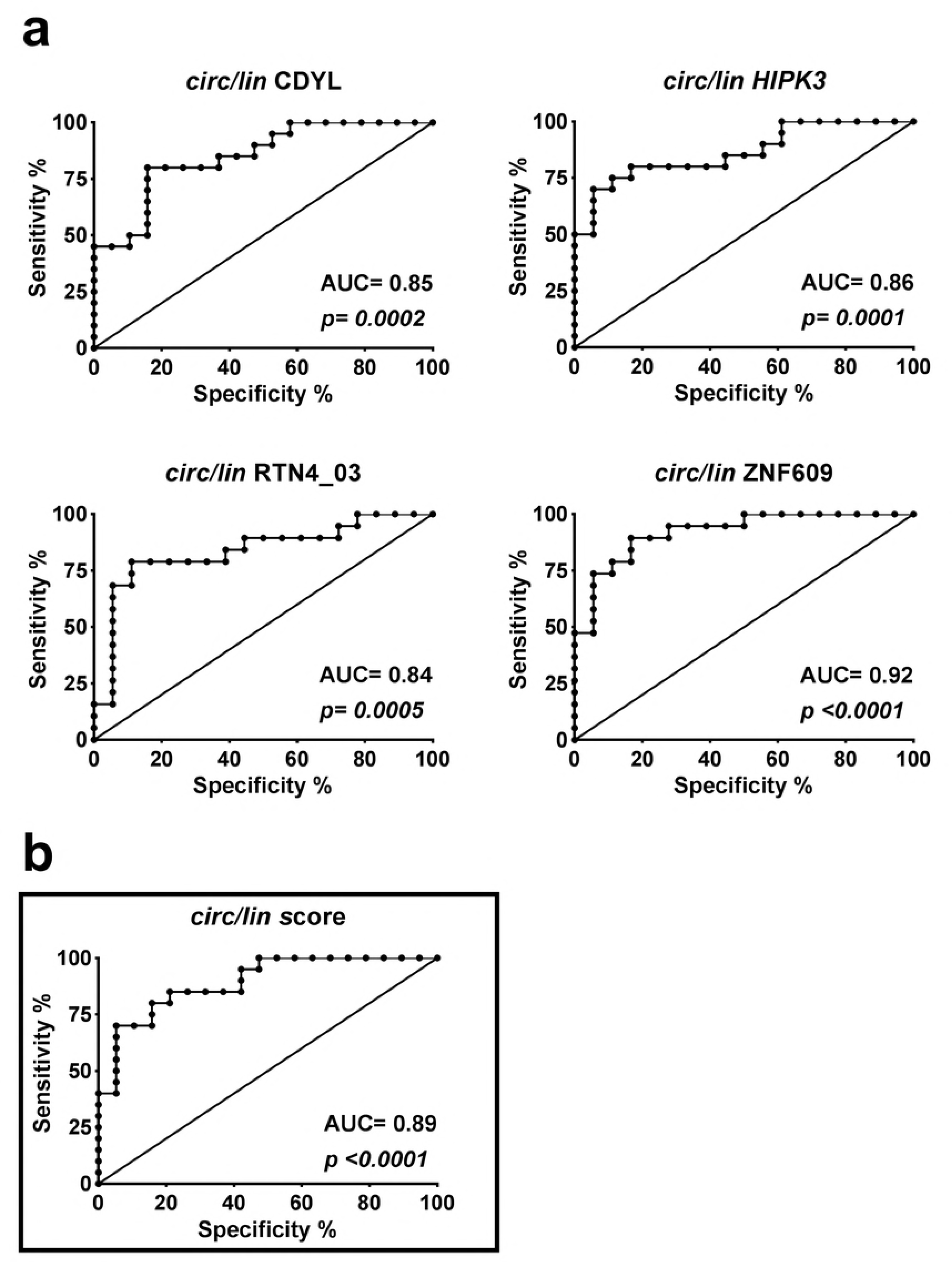
Discrimination of DM1 patients from controls using circular-to-linear ratios of DM1-circRNAs. **(a)** ROC curves show the sensitivity and specificity of each circRNA fraction (circ/lin) and of the combined “circular-to-linear score” to distinguish DM1 from healthy muscle tissue. **(b)**The “circular-to-linear score” was calculated by averaging the significantly modulated circular-to-linear ratios (circ/lin score, black rectangle). DM1= 20, CTRL= 19.

In conclusion, each of the five circular-to-linear ratios, as well as the combined “circular-to-linear score” of the DM1-circRNAs are useful to discriminate healthy form diseased patients.

### Correlation between DM1-circRNAs and clinical characteristics

To evaluate a potential relationship between the deregulation of circRNAs and clinical conditions, correlation analyses were performed. One of the most clinically relevant parameters for DM1 patients is muscle strength, measured by the Medical Research Council (MRC) grading system. We found that the changes of the circular fractions of circCDYL, circHIPK3, circRTN4_03 and circZNF609 displayed a significant negative correlation to MRC (Fig. 3a). Accordingly, a negative correlation was observed also between the MRC grading and the circular-to-linear score (Fig. 3b). The strongest and most significant correlation was found for the circular fraction of ZNF609, with Pearson r= 0.57 and p=0.0002.

**Figure 3.**
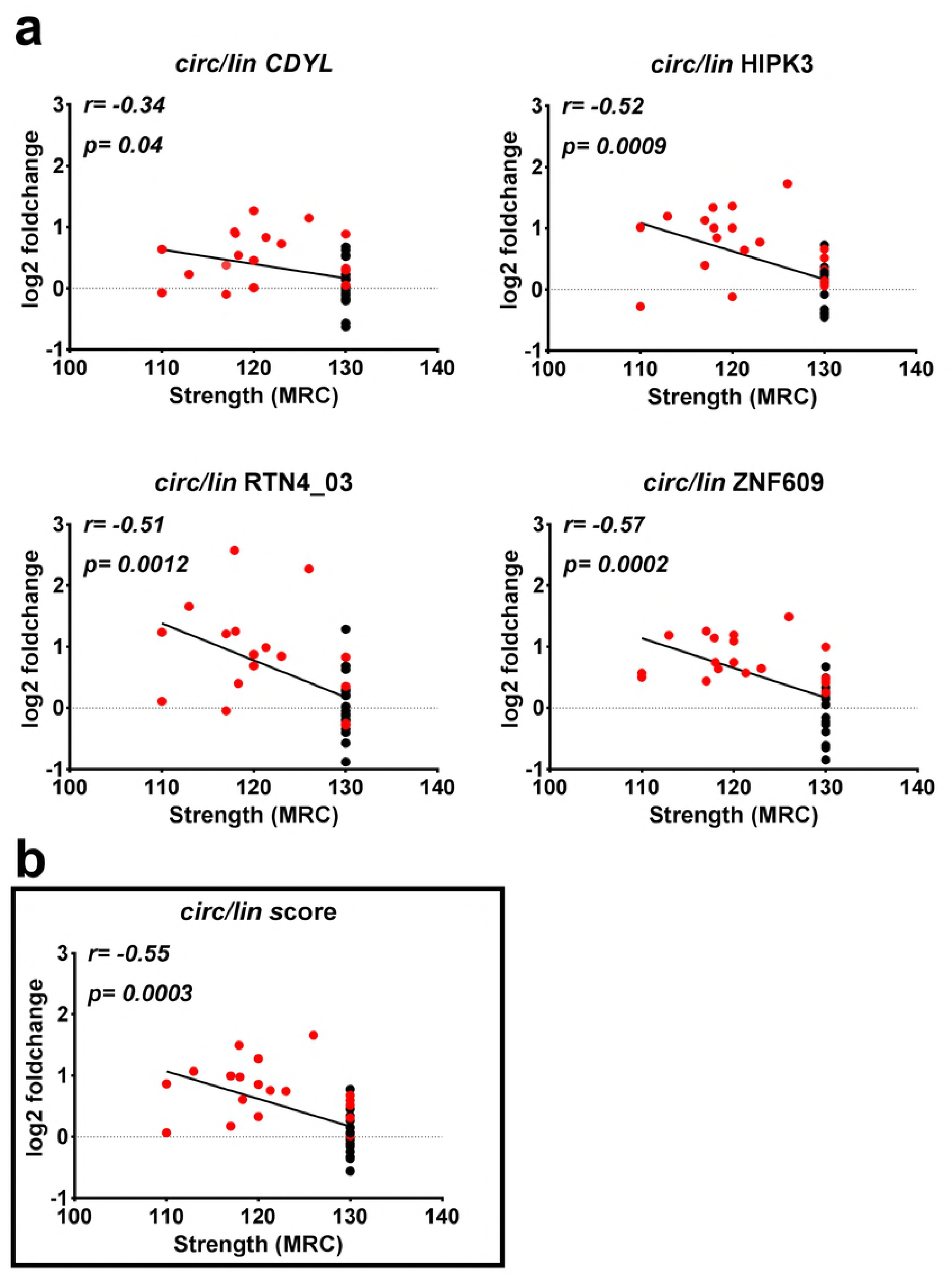
Correlation of muscle strength with circular-to-linear ratios of DM1-circRNAs. **(a)** Pearson’s correlation between significantly modulated circular-to-linear ratios identified in skeletal muscle biopsies and muscle strength measured by MRC megascore. **(b)** Pearson’s correlation of the “circular-to-linear score” (obtained averaging all DM1-circRNA fractions) and muscle strength measured by MRC megascore. DM1= 20 (red dots), CTRL= 19 (black dots).

Collectively, these data suggest a potential of circRNAs as DM1 biomarkers, in spite of the low number of subjects analyzed.

### DM1-circRNA levels in PBMCs and plasma of DM1 patients

Since peripheral blood can be obtained with a minimally invasive procedure, it represents a potentially interesting tissue for biomarker identification. Thus, we measured DM1-circRNA expression in PBMCs and plasma of DM1 patients and sex- and age-matched controls. Diseased and healthy subjects were chosen with the same criteria adopted for the harvesting of skeletal muscle biopsies (Table S4).

All DM1-circRNAs were readily detectable in PBMCs, but none of them showed a significant modulation (Fig. S4).

Among the DM1-circRNAs tested in plasma samples, circCDYL and circRTN4 were readily detectable. A small, but not significant induction could be observed in DM1 patients for circCDYL and circRTN4, consistent with the data obtained in biopsies (Fig. S5).

We conclude that the DM1-circRNA dysregulations observed in skeletal muscles are tissue restricted.

### DM1-circRNA expression in DM1 myogenic cell lines

We assessed whether circRNA alterations identified in DM1 muscle biopsies were also observed in cultured myoblasts. To this aim, we took advantage of DM1 and control muscle cell lines obtained by conversion of immortalized skin fibroblasts into multinucleated myotubes by forced expression of *MyoD1* [46]. In DM1 and control differentiated myogenic cells, all circRNAs tested were readily detectable. Interestingly, two of the circular transcripts, circRTN4 and circRTN4_03 were significantly increased in DM1 compared to control (Fig. S6). The analysis of their circular-to-linear ratio confirmed the induction of circRTN4_03. Additionally, an increased circular-to-linear ratio was also observed for ZNF609. These results are in line with the findings obtained in DM1 biopsies (Figure 4).

**Figure 4.**
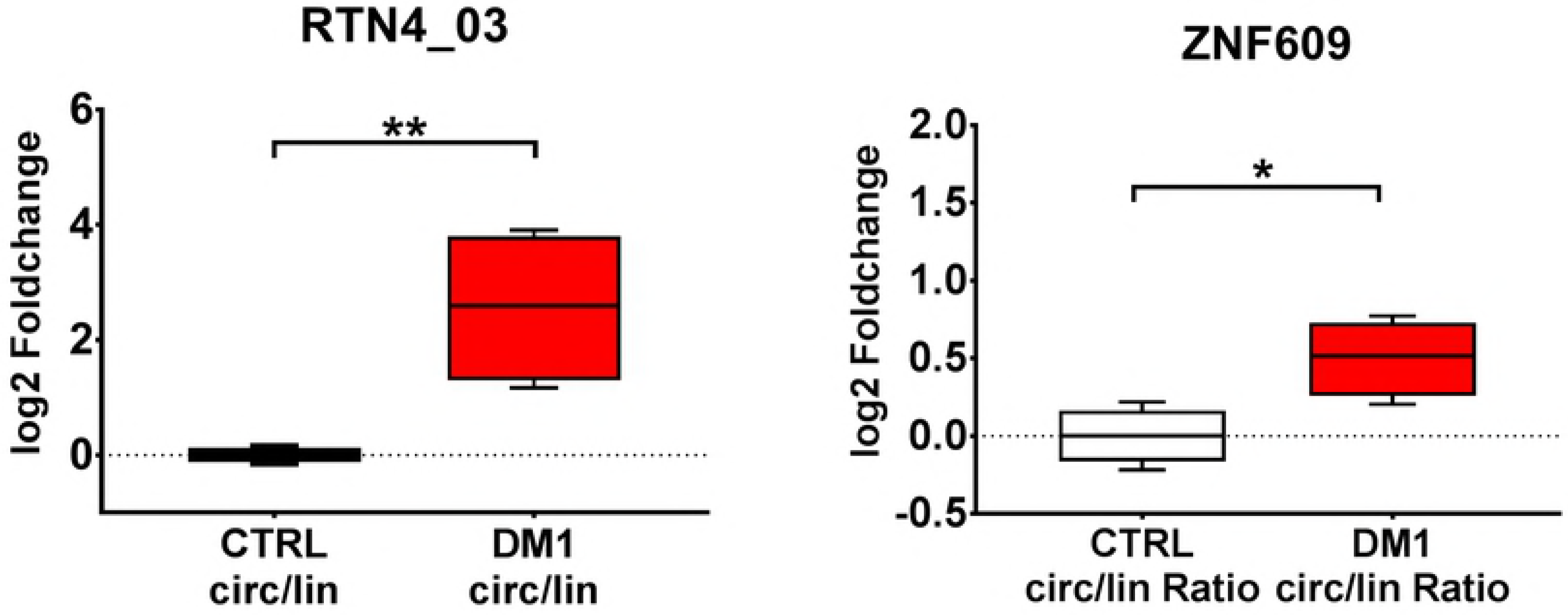
Differentially modulated circular-to-linear ratios in myogenic cell lines. Boxplots of significantly different circular-to-linear ratios identified by qPCR in differentiated DM1 myogenic cells compared to controls (*p<0.05; **p<0.01; DM1= 4; CTRL= 4).

We took advantage of this cell culture system to investigate whether silencing of DM1-related splicing factor affected the levels of the DM1-circRNAs modulated *in vitro*.

Since MBNL1 is impaired in DM1 patients [7, 9, 10], we assayed whether MBNL1 silencing in control myogenic cells induced, at least in part, the circRNA deregulations observed in DM1 myogenic cells. Control differentiated myogenic cells were transfected with MBNL1 siRNAs or relevant control oligonucleotides and RNA was extracted 3 days later. MBNL1 mRNA was significantly down modulated (Fig. S7a) and the expected alterations in the alternative-splicing patterns of SERCA1 and IR (Insulin Receptor) were observed (Fig. S7b and c) [50, 51]. However, no increase was observed in the abundance of circZNF609, circRTN4 and circRTN4_03 levels (Fig. S7d).

We also tested whether the silencing of CELF1, which is activated in patients and in DM1 disease models [52], rescued DM1-circRNA expression in DM1 differentiated myogenic cells. In spite of effective CELF1 knock-down (Fig. S8a), no significant change of circZNF609, circRTN and circRTN_03 levels was observed (Fig. S8b).

We conclude that the DM1 myogenic cell lines studied reflect, at least in part, the outcome of the DM1 biopsies and therefore represent a valuable tool for functional studies *in vitro*.

## Discussion

A molecular hallmark of DM1 is the dysregulation of alternative splicing, affecting many genes involved in muscle homeostasis and function [16, 17, 53]. CircRNAs are indeed alternative splicing products [20, 21] and, in this proof-of-principle study, we provide evidence of de-regulation of circRNA expression in DM1 patients. We analyzed 30 publicly available gene-expression data-sets of DM1 and control *tibialis anterior* muscles [36], using bioinformatics tools designed for the identification of circRNA-specific back-splice events. Applying stringent selection and abundance filters, we identified [1.800 unique circular splicing events, a number comparable to circRNAs found to be expressed in human myoblasts and myotubes [26].

During the testing phase of the bioinformatics pipeline with other data-sets, we performed several attempts of differential expression analysis (data not shown). Unfortunately, performances obtained were largely unsatisfactory, likely due to very low read numbers for most circRNA species and to normalization difficulties. Therefore, in this study we chose another approach for the identification of circRNAs potentially relevant in DM1. We filtered for circRNAs displaying expression across many samples and then selected circRNAs previously shown to be involved in myogenesis [26]. Of note, most of these circRNAs also displayed a high circular-to-linear ratio. This suggests that these circRNAs are not a mere by-product of the transcript maturation process, and might also indicate an independent regulation as well as an additional biological function.

For opportunity reasons, the validation step was performed in *biceps brachii* biopsies, since only for this muscle type a sufficient number of samples was available to us. It should be acknowledged that *biceps brachii* is generally less severely affected than other distal muscles in DM1 patients [1-3, 39]. Thus, some of the circRNA level differences that failed to reach statistical significance in our validation analysis, might be indeed relevant in distal muscles. On the other side, it is plausible to hypothesize that circRNA alterations identified in proximal muscles could be more pronounced in distal muscles.

In spite of the preliminary and not-comprehensive nature of this study, we found that the levels of 5 out of 9 circRNAs tested were significantly increased, suggesting a potentially pervasive dysregulation of circRNAs in DM1. Accordingly, 4 of these circRNAs also displayed an increased circular-to-linear ratio. The most likely interpretation of these alterations is that they are caused by the dysfunction of the alternative splicing machinery characterizing DM1 [16, 17, 53]. Silencing of either MBNL1 or CELF1 in control and DM1 cultured myotubes did not affect the abundance of circZNF609, circRTN4 and circRTN4_03. While negative data should always be evaluated in a very cautious manner, one possible interpretation is that other splicing factors, such as different MBNL-family members, Staufen1 and DDX5 [6-12], regulate the generation of the DM1-circRNAs. Moreover, these splicing factors might be redundant in circRNA regulation, making the silencing of a single splicing factor ineffective. Finally, higher stability of circRNAs compared to their linear counterparts [54] should also be considered as possible mechanism underpinning the increase of the circular-to-linear ratio of DM1-circRNAs.

CircRNA dysregulation may lead to pathological consequences to be investigated. In this respect, however, little is known of the identified DM1-circRNAs. CircHIPK3 positively regulates human cell growth by sponging multiple miRNAs [55, 56]. Among these miRNAs, there are miR-29b and miR-193a. Intriguingly, both miRNAs were previously found to be down-modulated in DM1 [57] and DM2, respectively [47]. CircHIPK3 levels are also increased in retinal endothelial cells exposed to diabetes-related stressors, in retinas of diabetic mice and in the plasma of diabetic patients. Moreover, in retinal endothelial-cells, circHIPK3 affects cell viability, proliferation, migration, and function [58]. While the implication of circHIPK3 in DM1 should be investigated, it is worth noting that insulin resistance is very often present in DM1 patients [1-3, 59].

CircZNF609 is downregulated during myogenesis and can specifically control myoblast proliferation [26]. Moreover, its mouse homologue circZfp609 suppresses myogenic differentiation [60]. Of note, circZNF609 is elevated in Duchenne muscular dystrophy myoblasts, indicating that the DM1-circRNAs identified in this study might be deregulated in other muscular diseases. Like circHIPK3, also circZNF609 is induced by high glucose *in vivo* and *in vitro* and regulates the function of retinal endothelial cells [61].

Finally, the host gene of circRTN4 displays muscle-specific splicing [62]. It is a direct target of RBM20, an alternative-splicing regulator of cardiac genes, associated with coronary heart disease. Mutation of RBM20 results in altered splicing of its target genes, causing the retention of specific exons of RTN4 mRNA. Since RBM20 is also expressed and active in skeletal muscles [63], further investigations are needed to assess the potential involvement of RBM20 also in circRTN4 formation.

Ashwal-Fluss and collaborators [33], by studying circRNAs identified in neuronal tissues, reported that the second exon of the splicing factor *MBLN1* (*MBL* in drosophila) is circularized in flies and humans and that circMBL production competes with canonical pre-mRNA splicing. Moreover, Muscleblind protein interacts with flanking introns of its own gene to promote exon circularization. This observation prompted us to measure circMBNL1 in skeletal muscle biopsies and PBMCs of DM1 patients, where MBNL1 protein bioavailability is reduced by its sequestration in nuclear CUG-foci [9, 10]. However, we did not observe any modulation of circMBNL1 in these tissues. The most likely explanation is that circMBNL1 regulation is highly context specific. Indeed, while in fly heads, this circRNA is more abundant than the linear counterpart, the opposite seems to be true in human skeletal muscle, where RNAseq data indicated a circular-to-linear ratio of 0.05 (Table S3). In keeping with this hypothesis, very low circMBL levels were also observed in drosophila S2 cells [33]. Furthermore, changes in the splicing pattern of MBNL1 mRNA (comprising or not exon1) were observed in cardiac and skeletal muscles, but not in the brain of DM1 patients compared to controls [64, 65], confirming a high tissue-specificity in the regulation of MBNL1 transcript.

Identification of diagnostic and prognostic biomarkers is an unmet clinical need for DM1 patients. Recent studies suggest that alternative splicing isoforms in skeletal muscle tissue have a high potential as biomarkers of DM severity and for the monitoring of therapeutic responses [53]. Due to their loop-structure, circRNAs are highly resistant to exonucleases [54], holding a great potential as disease biomarkers. Promisingly, the potential of circRNAs as molecular markers has been highlighted recently in various types of cancer [66], measuring circRNAs not only in biopsies of solid tissues, but also in extracellular compartments, such as serum or exosomes [66, 67].

It is still too early to conclude whether circRNAs will find their way to the clinic. This will largely depend on both technical issues, such as detectability and stability of circRNAs in biological samples, as well as on whether circRNAs are functional elements of the molecular mechanisms driving the disease or mere byproducts [54]. While further studies are obviously needed, we found that the circular-to-linear ratios as well as the combined “circular-to-linear score” of the DM1-circRNAs in skeletal muscle biopsies accurately discriminated healthy form DM1 patients. Moreover, a correlation between the MRC grading and the circular-to-linear score could be identified. Additional investigations are needed to evaluate the potential of circRNAs as DM1 biomarker, involving a higher number of subjects. Comparison with other myopathies will also allow to investigate their possible disease-specificity.

CircRNAs are detectable also in the peripheral blood that can be harvested with minimally-invasive techniques. Indeed, DM1-circRNAs were measured both in PBMCs and, at least in part, in plasma samples. On the one side, the high context-dependent regulation of circRNAs implicates that the pattern of expression observed in the peripheral blood is unlikely to mirror that of other tissues, such as the skeletal muscle. On the other side, the apparent pervasiveness of circRNA dysregulation suggest that comprehensive screenings of circRNAs might indeed succeed in the identification of common circRNAs between blood and affected organs. An alternative approach may be represented by the analysis of urine extracellular RNAs, whose splice variants have been shown to discriminate DM1 patients efficiently [68].

## Acknowledgments

We thank the platform for immortalization of human cells of the Institut de Myologie, Paris, France, for providing DM1 and control myogenic cell lines and Michele Cavalli (Biomedical Sciences for Health, University of Milan and IRCCS Policlinico San Donato, Milan, Italy) for his support in clinical data collection.

## Supporting information

**Figure S1. Histopathological analysis of *biceps brachii* biopsies obtained from a representative DM1 patient (a, b) and from a representative control (c). In DM1 patient, Hematoxylin & Eosin (a) and ATPase pH 10.4 (b) stainings displayed the characteristic histopathological features of DM1,** such as central nuclei (asterisks), atrophic fibers (arrow) and an evident fiber size variability both of type 1 (negative fibers) and type 2 (brown fibers) fibers. **Figure S2. Significantly modulated circRNAs in DM1 skeletal muscles.** Scatterplots of circRNA (circ.) transcripts identified by qPCR as differentially expressed in DM1 *biceps brachii* biopsies are shown together with their linear counterparts (lin.). Lines indicate mean and standard error values for each group. After normality test, statistical significance was calculated either by t-test or Mann-Whitney test (threshold p<0.05), followed by correction multiple comparison, with significance threshold set at q<0.01 (*q<0.01; **q<0.001). DM1= 20 (red dots), Controls= 19 (CTRL, black dots).

**Figure S3. Discrimination of DM1 from healthy patients by circRNA modulation.** ROC curves show the sensitivity and specificity of each DM1-circRNA to distinguish DM1 from healthy muscle tissue. DM1= 20, CTRL= 19.

**Figure S4. DM1-circRNA levels in PBMCs derived from DM1 patients and controls. RNA was** extracted from PBMCs of DM1 and control individuals. Scatterplots show the levels of DM1-circRNAs detected by qPCR. DM1= 19 (red dots), Controls= 18 (CTRL, black dots). Lines indicate mean and standard error values for each group.

**Figure S5. DM1-circRNA levels in plasma derived from DM1 patients and controls. RNA was** extracted from platelet-free plasma of DM1 and control individuals. Scatterplots show the levels of circRNAs detectable by qPCR. DM1= 29 (red dots), Controls= 28 (CTRL, black dots). Lines indicate mean and standard error values for each group.

**Figure S6. Significantly modulated circRNAs in DM1 myogenic cell lines.** Boxplots of differentially expressed circRNAs (circ.) and their linear counterparts (lin.) identified by qPCR in differentiated DM1 myogenic cells compared to controls (*p<0.05; **p<0.01; DM1= 4; CTRL= 4).

**Figure S7. MBNL1 silencing does not affect DM1-circRNA levels. Differentiated control** myogenic cells were transfected with siRNAs targeting MBNL1 or with control siRNAs (n=5). (**a**) Efficiency of MBNL1 knock-down was assessed by qPCR (*** p<0.0001). (**b** and **c**) MBNL1 silencing induced the expected increases of the SERCA1 isoform excluding exon 22 (isoform b), and of the IR isoform excluding exon 11 (isoform a), as assessed by PCR followed by agarose gel electrophoresis. Representative gels are shown. (**d**) circZNF609, circRTN4 and circRTN4_03 levels were measured by qPCR. None of the circRNAs displayed a statistically significant increase.

**Figure S8. CELF1 silencing does not affect DM1-circRNA levels. Differentiated DM1 myogenic** cells were transfected with siRNAs targeting CELF1 or with control siRNAs (n=3). (**a**) Efficiency of CELF1 knock-down was assessed by qPCR (*** p<0.0001). (**b**) circZNF609, circRTN4 and circRTN4_03 levels were measured by qPCR. None of the circRNAs displayed a statistically significant decrease.

**Table S1.** Library size of publicly available data-sets used for circRNA identification in DM1 **skeletal muscle by RNAseq**. A set of 30 transcriptomes (25 DM1 and 5 healthy controls) from human tibialis biopsies (GSE86356) was investigated for back-splice events. The sequencing depth is reported as million sequenced reads after mapping to the human genome version hg19. Each data-set is identified by its SRA Run number; controls are highlighted in grey.

**Table S2. circRNA identification and expression in DM1 skeletal muscle by RNAseq.** Normalized counts of 1797 different circRNA species identified by CIRI2-algorithm, following a filtering step for abundance. Each back-splice junction is identified with the coordinates of the involved donor and acceptor sites in the format “chromosome number: donor position | acceptor position”. Each data-set is identified by its SRA Run number, controls are highlighted in grey.

**Table S3. Ratios of circular versus linear expression levels measured in DM1 skeletal muscle by RNAseq.** For estimation of circular-to-linear ratios, the linear junction with the highest coverage involved with either the donor or the acceptor site of the back-splice event was determined. The averaged, normalized counts across all libraries in each condition were calculated for linear and back-splice junction. The circular-to-linear ratios were determined for controls and DM1 and are here highlighted in grey. Additionally, the circRNAs identified in human tibialis biopsies were intersected with myogenic circRNAs identified by Legnini et al. [26] The final validation set chosen for qPCR is displayed in the column “Validation-set”.

**Table S4. Clinical data of DM1 patients and controls used for circRNA detection in plasma and PBMCs.** NR: not relevant, NA: not available.

**Table S5. Primer-sequences. List of primer-couples used for relative quantification of circRNAs** and their linear counterparts by qPCR. With the exception of CDYL and HIPK3, circular transcripts and their linear counterparts shared one primer, either forward or reverse.

## References

1 Meola G, Cardani R. Myotonic dystrophies: An update on clinical aspects, genetic, pathology, and molecular pathomechanisms. Biochim Biophys Acta. 2015;1852: 594–606.

2 De Antonio M, Dogan C, Hamroun D, Mati M, Zerrouki S, Eymard B, et al. Unravelling the myotonic dystrophy type 1 clinical spectrum: A systematic registry-based study with implications for disease classification. Rev Neurol (Paris). 2016;172: 572–580.

3 Gagnon C, Chouinard MC, Laberge L, Veillette S, Begin P, Breton R, et al. Health supervision and anticipatory guidance in adult myotonic dystrophy type 1. Neuromuscul Disord. 2010;20: 847-851.

4 Gourdon G, Meola G. Myotonic Dystrophies: State of the Art of New Therapeutic Developments for the CNS. Front Cell Neurosci. 2017;11: 101.

5 Fu YH, Pizzuti A, Fenwick RG, Jr, King J, Rajnarayan S, Dunne PW, et al. An unstable triplet repeat in a gene related to myotonic muscular dystrophy. Science. 1992;255: 1256–1258.

6 Bondy-Chorney E, Crawford Parks TE, Ravel-Chapuis A, Klinck R, Rocheleau L, Pelchat M, et al. Staufen1 Regulates Multiple Alternative Splicing Events either Positively or Negatively in DM1 Indicating Its Role as a Disease Modifier. PLoS Genet. 2016;12: e1005827.

7 Lin X, Miller JW, Mankodi A, Kanadia RN, Yuan Y, Moxley RT, et al. Failure of MBNL1-dependent post-natal splicing transitions in myotonic dystrophy. Hum Mol Genet. 2006;15: 2087-2097.

8 Laurent FX, Sureau A, Klein AF, Trouslard F, Gasnier E, Furling D, et al. New function for the RNA helicase p68/DDX5 as a modifier of MBNL1 activity on expanded CUG repeats. Nucleic Acids Res. 2012;40: 3159–3171.

9 Dansithong W, Paul S, Comai L, Reddy S. MBNL1 is the primary determinant of focus formation and aberrant insulin receptor splicing in DM1. J Biol Chem. 2005;280: 5773–5780.

10 Miller JW, Urbinati CR, Teng-Umnuay P, Stenberg MG, Byrne BJ, Thornton CA, et al. Recruitment of human muscleblind proteins to (CUG)(n) expansions associated with myotonic dystrophy. EMBO J. 2000;19: 4439–4448.

11 Philips AV, Timchenko LT, Cooper TA. Disruption of splicing regulated by a CUG-binding protein in myotonic dystrophy. Science. 1998;280: 737–741.

12 Timchenko LT, Miller JW, Timchenko NA, DeVore DR, Datar KV, Lin L, et al. Identification of a (CUG)n triplet repeat RNA-binding protein and its expression in myotonic dystrophy. Nucleic Acids Res. 1996;24: 4407–4414.

13 Chen KY, Pan H, Lin MJ, Li YY, Wang LC, Wu YC, et al. Length-dependent toxicity of untranslated CUG repeats on Caenorhabditis elegans. Biochem Biophys Res Commun. 2007;352: 774–779.

14 Garcia-Lopez A, Monferrer L, Garcia-Alcover I, Vicente-Crespo M, Alvarez-Abril MC, Artero RD. Genetic and chemical modifiers of a CUG toxicity model in Drosophila. PLoS One. 2008;3: e1595.

15 Mankodi A, Logigian E, Callahan L, McClain C, White R, Henderson D, et al. Myotonic dystrophy in transgenic mice expressing an expanded CUG repeat. Science. 2000;289: 1769–1773.

16 Du H, Cline MS, Osborne RJ, Tuttle DL, Clark TA, Donohue JP, et al. Aberrant alternative splicing and extracellular matrix gene expression in mouse models of myotonic dystrophy. Nat Struct Mol Biol. 2010;17: 187–193.

17 Kalsotra A, Xiao X, Ward AJ, Castle JC, Johnson JM, Burge CB, et al. A postnatal switch of CELF and MBNL proteins reprograms alternative splicing in the developing heart. Proc Natl Acad Sci U S A. 2008;105: 20333–20338.

18 Kanadia RN, Johnstone KA, Mankodi A, Lungu C, Thornton CA, Esson D, et al. A muscleblind knockout model for myotonic dystrophy. Science. 2003;302: 1978–1980.

19 Batra R, Charizanis K, Manchanda M, Mohan A, Li M, Finn DJ, et al. Loss of MBNL leads to disruption of developmentally regulated alternative polyadenylation in RNA-mediated disease. Mol Cell. 2014;56: 311–322.

20 Noh JH, Kim KM, McClusky WG, Abdelmohsen K, Gorospe M. Cytoplasmic functions of long noncoding RNAs. Wiley Interdiscip Rev RNA. 2018;9: e1471.

21 Holdt LM, Kohlmaier A, Teupser D. Molecular roles and function of circular RNAs in eukaryotic cells. Cell Mol Life Sci. 2018;75: 1071–1098.

22 Du WW, Zhang C, Yang W, Yong T, Awan FM, Yang BB. Identifying and Characterizing circRNA-Protein Interaction. Theranostics. 2017;7: 4183–4191.

23 Carrara M, Fuschi P, Ivan C, Martelli F. Circular RNAs: Methodological challenges and perspectives in cardiovascular diseases. J Cell Mol Med. 2018.

24 Jeck WR, Sorrentino JA, Wang K, Slevin MK, Burd CE, Liu J, et al. Circular RNAs are abundant, conserved, and associated with ALU repeats. RNA. 2013;19: 141–157.

25 Memczak S, Jens M, Elefsinioti A, Torti F, Krueger J, Rybak A, et al. Circular RNAs are a large class of animal RNAs with regulatory potency. Nature. 2013;495: 333–338.

26 Legnini I, Di Timoteo G, Rossi F, Morlando M, Briganti F, Sthandier O, et al. Circ-ZNF609 Is a Circular RNA that Can Be Translated and Functions in Myogenesis. Mol Cell. 2017;66: 22–37.e9.

27 Salzman J, Gawad C, Wang PL, Lacayo N, Brown PO. Circular RNAs are the predominant transcript isoform from hundreds of human genes in diverse cell types. PLoS One. 2012;7: e30733.

28 Rybak-Wolf A, Stottmeister C, Glazar P, Jens M, Pino N, Giusti S, et al. Circular RNAs in the Mammalian Brain Are Highly Abundant, Conserved, and Dynamically Expressed. Mol Cell. 2015;58: 870–885.

29 Wei X, Li H, Yang J, Hao D, Dong D, Huang Y, et al. Circular RNA profiling reveals an abundant circLMO7 that regulates myoblasts differentiation and survival by sponging miR-378a-3p. Cell Death Dis. 2017;8: e3153.

30 Hansen TB, Jensen TI, Clausen BH, Bramsen JB, Finsen B, Damgaard CK, et al. Natural RNA circles function as efficient microRNA sponges. Nature. 2013;495: 384–388.

31 Piwecka M, Glazar P, Hernandez-Miranda LR, Memczak S, Wolf SA, Rybak-Wolf A, et al. Loss of a mammalian circular RNA locus causes miRNA deregulation and affects brain function. Science. 2017;357: 10.1126/science.aam8526. Epub 2017 Aug 10.

32 Li Z, Huang C, Bao C, Chen L, Lin M, Wang X, et al. Exon-intron circular RNAs regulate transcription in the nucleus. Nat Struct Mol Biol. 2015;22: 256–264.

33 Ashwal-Fluss R, Meyer M, Pamudurti NR, Ivanov A, Bartok O, Hanan M, et al. circRNA biogenesis competes with pre-mRNA splicing. Mol Cell. 2014;56: 55–66.

34 Pamudurti NR, Bartok O, Jens M, Ashwal-Fluss R, Stottmeister C, Ruhe L, et al. Translation of CircRNAs. Mol Cell. 2017;66: 9–21.e7.

35 Yang Z, Xie L, Han L, Qu X, Yang Y, Zhang Y, et al. Circular RNAs: Regulators of Cancer-Related Signaling Pathways and Potential Diagnostic Biomarkers for Human Cancers. Theranostics. 2017;7: 3106–3117.

36 Wagner SD, Struck AJ, Gupta R, Farnsworth DR, Mahady AE, Eichinger K, et al. Dose-Dependent Regulation of Alternative Splicing by MBNL Proteins Reveals Biomarkers for Myotonic Dystrophy. PLoS Genet. 2016;12: e1006316.

37 Gao Y, Wang J, Zhao F. CIRI: an efficient and unbiased algorithm for de novo circular RNA identification. Genome Biol. 2015;16: 4-014–0571-3.

38 Gao Y, Zhang J, Zhao F. Circular RNA identification based on multiple seed matching. Brief Bioinform. 2017.

39 Udd B, Meola G, Krahe R, Thornton C, Ranum LP, Bassez G, et al. 140th ENMC International Workshop: Myotonic Dystrophy DM2/PROMM and other myotonic dystrophies with guidelines on management. Neuromuscul Disord. 2006;16: 403–413.

40 Valaperta R, Sansone V, Lombardi F, Verdelli C, Colombo A, Valisi M, et al. Identification and characterization of DM1 patients by a new diagnostic certified assay: neuromuscular and cardiac assessments. Biomed Res Int. 2013;2013: 958510.

41 Mathieu J, Boivin H, Meunier D, Gaudreault M, Begin P. Assessment of a disease-specific muscular impairment rating scale in myotonic dystrophy. Neurology. 2001;56: 336–340.

42 Voellenkle C, van Rooij J, Cappuzzello C, Greco S, Arcelli D, Di Vito L, et al. MicroRNA signatures in peripheral blood mononuclear cells of chronic heart failure patients. Physiol Genomics. 2010;42: 420–426.

43 Perfetti A, Greco S, Bugiardini E, Cardani R, Gaia P, Gaetano C, et al. Plasma microRNAs as biomarkers for myotonic dystrophy type 1. Neuromuscul Disord. 2014;24: 509–515.

44 Perfetti A, Greco S, Cardani R, Fossati B, Cuomo G, Valaperta R, et al. Validation of plasma microRNAs as biomarkers for myotonic dystrophy type 1. Sci Rep. 2016;6: 38174.

45 Dubowitz V. Muscle biopsy: A Practical Approach. In: Anonymous Muscle biopsy: A Practical Approach. London: Bailliere Tindall: Dubowitz, Victor; 1985. pp. 19–40.

46 Arandel L, Polay Espinoza M, Matloka M, Bazinet A, De Dea Diniz D, Naouar N, et al. Immortalized human myotonic dystrophy muscle cell lines to assess therapeutic compounds. Dis Model Mech. 2017;10: 487–497.

47 Greco S, Perfetti A, Fasanaro P, Cardani R, Capogrossi MC, Meola G, et al. Deregulated microRNAs in myotonic dystrophy type 2. PLoS One. 2012;7: e39732.

48 Greco S, De Simone M, Colussi C, Zaccagnini G, Fasanaro P, Pescatori M, et al. Common micro-RNA signature in skeletal muscle damage and regeneration induced by Duchenne muscular dystrophy and acute ischemia. FASEB J. 2009;23: 3335–3346.

49 Livak KJ, Schmittgen TD. Analysis of relative gene expression data using real-time quantitative PCR and the 2(-Delta Delta C(T)) Method. Methods. 2001;25: 402–408.

50 Kimura T, Nakamori M, Lueck JD, Pouliquin P, Aoike F, Fujimura H, et al. Altered mRNA splicing of the skeletal muscle ryanodine receptor and sarcoplasmic/endoplasmic reticulum Ca2+-ATPase in myotonic dystrophy type 1. Hum Mol Genet. 2005;14: 2189–2200.

51 Savkur RS, Philips AV, Cooper TA. Aberrant regulation of insulin receptor alternative splicing is associated with insulin resistance in myotonic dystrophy. Nat Genet. 2001;29: 40–47.

52 Wang ET, Ward AJ, Cherone JM, Giudice J, Wang TT, Treacy DJ, et al. Antagonistic regulation of mRNA expression and splicing by CELF and MBNL proteins. Genome Res. 2015;25: 858–871.

53 Nakamori M, Sobczak K, Puwanant A, Welle S, Eichinger K, Pandya S, et al. Splicing biomarkers of disease severity in myotonic dystrophy. Ann Neurol. 2013;74: 862–872.

54 Ebbesen KK, Kjems J, Hansen TB. Circular RNAs: Identification, biogenesis and function. Biochim Biophys Acta. 2016;1859: 163–168.

55 Zheng Q, Bao C, Guo W, Li S, Chen J, Chen B, et al. Circular RNA profiling reveals an abundant circHIPK3 that regulates cell growth by sponging multiple miRNAs. Nat Commun. 2016;7: 11215.

56 Liu X, Liu B, Zhou M, Fan F, Yu M, Gao C, et al. Circular RNA HIPK3 regulates human lens epithelial cells proliferation and apoptosis by targeting the miR-193a/CRYAA axis. Biochem Biophys Res Commun. 2018;503: 2277–2285.

57 Perbellini R, Greco S, Sarra-Ferraris G, Cardani R, Capogrossi MC, Meola G, et al. Dysregulation and cellular mislocalization of specific miRNAs in myotonic dystrophy type 1. Neuromuscul Disord. 2011;21: 81–88.

58 Shan K, Liu C, Liu BH, Chen X, Dong R, Liu X, et al. Circular Noncoding RNA HIPK3 Mediates Retinal Vascular Dysfunction in Diabetes Mellitus. Circulation. 2017;136: 1629–1642.

59 Renna LV, Bose F, Iachettini S, Fossati B, Saraceno L, Milani V, et al. Receptor and post-receptor abnormalities contribute to insulin resistance in myotonic dystrophy type 1 and type 2 skeletal muscle. PLoS One. 2017;12: e0184987.

60 Wang Y, Li M, Wang Y, Liu J, Zhang M, Fang X, et al. A Zfp609 circular RNA regulates myoblast differentiation by sponging miR-194–5p. Int J Biol Macromol. 2018.

61 Liu C, Yao MD, Li CP, Shan K, Yang H, Wang JJ, et al. Silencing Of Circular RNA-ZNF609 Ameliorates Vascular Endothelial Dysfunction. Theranostics. 2017;7: 2863–2877.

62 Maatz H, Jens M, Liss M, Schafer S, Heinig M, Kirchner M, et al. RNA-binding protein RBM20 represses splicing to orchestrate cardiac pre-mRNA processing. J Clin Invest. 2014;124: 3419–3430.

63 Chen Z, Maimaiti R, Zhu C, Cai H, Stern A, Mozdziak P, et al. Z-band and M-band titin splicing and regulation by RNA binding motif 20 in striated muscles. J Cell Biochem. 2018.

64 Konieczny P, Stepniak-Konieczna E, Sobczak K. MBNL proteins and their target RNAs, interaction and splicing regulation. Nucleic Acids Res. 2014;42: 10873–10887.

65 Konieczny P, Stepniak-Konieczna E, Taylor K, Sznajder LJ, Sobczak K. Autoregulation of MBNL1 function by exon 1 exclusion from MBNL1 transcript. Nucleic Acids Res. 2017;45: 1760-1775.

66 Kristensen LS, Hansen TB, Veno MT, Kjems J. Circular RNAs in cancer: opportunities and challenges in the field. Oncogene. 2018;37: 555–565.

67 Li Y, Zheng Q, Bao C, Li S, Guo W, Zhao J, et al. Circular RNA is enriched and stable in exosomes: a promising biomarker for cancer diagnosis. Cell Res. 2015;25: 981–984.

68 Antoury L, Hu N, Balaj L, Das S, Georghiou S, Darras B, et al. Analysis of extracellular mRNA in human urine reveals splice variant biomarkers of muscular dystrophies. Nat Commun. 2018;9: 3906-018–06206-0.

